# Caspase-4 dimerisation and D289 auto-processing elicits an interleukin-1β converting enzyme

**DOI:** 10.1101/2023.01.05.522955

**Authors:** Amy H. Chan, Kassandra Vezyrgiannis, Jessica B. Von Pein, Xiaohui Wang, Larisa I Labzin, Dave Boucher, Kate Schroder

**Author notes:** Equal contribution (Dave Boucher, Kate Schroder).

## Abstract

The non-canonical inflammasome is a signalling complex critical for cell defence against cytosolic Gram-negative bacteria. A key step in the human non-canonical inflammasome pathway involves unleashing the proteolytic activity of caspase-4 within this complex. Caspase-4 induces inflammatory responses by cleaving gasdermin-D (GSDMD) to initiate pyroptosis, although the molecular mechanisms that activate caspase-4 and govern its capacity to cleave substrates are poorly defined. Caspase-11, the murine counterpart of caspase-4, acquires protease activity within the non-canonical inflammasome by forming a dimer that self-cleaves at D285 to directly cleave GSDMD. These cleavage events trigger signalling via the NLRP3-ASC-caspase-1 axis, leading to downstream cleavage of the pro-interleukin-1β (pro-IL-1β) cytokine precursor. Here, we show that caspase-4 first dimerises then self-cleaves at two sites – D270 and D289 – in the interdomain linker to acquire full proteolytic activity, cleave GSDMD and induce cell death. Surprisingly, caspase-4 dimerisation and self-cleavage at D289 generates a caspase-4 p34/p9 protease species that directly cleaves pro-IL-1β, resulting in its maturation and secretion independently of the NLRP3 inflammasome in primary human myeloid and epithelial cells. Our study thus elucidates the key molecular events that underpin signalling by the caspase-4 inflammasome, and identifies IL-1β as a natural substrate of caspase-4.

## Introduction

Non-canonical inflammasome activity is critical for innate immune responses to lipopolysaccharide (LPS) on cytosolic Gram-negative bacteria (Kayagaki *et al*, 2011; Kayagaki *et al*, 2013; Shi *et al*, 2014). In mice, the non-canonical inflammasome activates the cysteine protease, caspase-11, while canonical inflammasomes activate caspase-1 (Ross *et al*, 2022). Caspase-4 and -5 are the human orthologs of murine caspase-11 and are presumed to be activated by similar mechanisms and exert similar biological functions. In line with this, all three caspases of non-canonical inflammasomes (hereafter called caspase-4/5/11) detect and interact with LPS with the assistance of guanylate-binding proteins (GBPs) (Fisch *et al*, 2019; Kutsch *et al*, 2020; Santos *et al*, 2020; Wandel *et al*, 2020). LPS-activated caspase-4/5/11 subsequently cleaves the substrate gasdermin-D (GSDMD) to generate a GSDMD p30 fragment that inserts into the plasma membrane and forms pores. GSDMD pores trigger either ninjurin-1-dependent plasma membrane rupture and pyroptotic cell death (Aglietti *et al*, 2016; Ding *et al*, 2016; He *et al*, 2015; Kayagaki *et al*, 2021; Kayagaki *et al*, 2015; Liu *et al*, 2016; Shi *et al*, 2015) or pyroptosis-associated extrusion of neutrophil extracellular traps (Chen *et al*, 2018). GSDMD pores also trigger ionic flux and the resultant assembly of NLRP3-ASC-caspase-1 inflammasomes, which in turn generate active caspase-1 that cleaves pro-IL-1β and pro-IL-18 into their mature forms (Boucher *et al*, 2018; Chan & Schroder, 2019; Monteleone *et al*, 2018; Rühl & Broz, 2015).

While the mechanisms controlling caspase-1 and -11 activation are now defined (Boucher *et al*., 2018; Ross *et al*, 2018), those controlling human caspase-4/5 remain poorly understood. It is likely that dimerisation is required for the activation of caspase-4, as this underpins the activation of related initiator caspases (caspase-1, -8, -9, -11) (Boatright *et al*, 2003; Boucher *et al*., 2018; Ross *et al*., 2018). Caspase-1, -8 and -9 are synthesised as inactive monomeric zymogens composed of a catalytic domain (large plus small subunits of the protease) connected to an N-terminal domain (CARD or DED). The CARD domain enables recruitment to signalling platforms (Martinon *et al*, 2002; Pop *et al*, 2006) where clustering facilitates proximity-induced dimerisation of the caspase-1/8/9 enzymatic subunits to enable caspase intrinsic proteolytic activity (Boatright *et al*., 2003; Boucher *et al*., 2018; Renatus *et al*, 2001). Specifically, dimerisation induces autoproteolysis within either the linker that connects the protease to its N-terminal recruitment domain (e.g. the caspase-1 CARD-domain linker; CDL), or the linker that connects the two catalytic subunits (interdomain linker; IDL). The impact of IDL or CDL processing on protease activity and substrate repertoire varies depending on the caspase (Boucher *et al*., 2018; Oberst *et al*, 2010; Pop *et al*., 2006). IDL processing broadens the caspase-1 and -8 substrate repertoire, while CDL processing deactivates caspase-1 by ejecting dimers from the inflammasome. Dimer detachment destabilises caspase-1, causing it to dissociate into inactive monomers (Boucher *et al*., 2018), while IDL cleavage is necessary for caspase-1 to cleave cytokines. Interestingly, IDL cleavage may be dispensable for caspase-1-induced death of murine cells (Broz *et al*, 2010; Dick *et al*, 2016), suggesting that auto-processing is not always necessary for specific functions of inflammatory caspases.

Caspase-4 is the closest orthologue of murine caspase-11 (Baker *et al*, 2015; Casson *et al*, 2015), and LPS is reported as a direct ligand of caspase-4 and -11. Indeed, binding of their CARD domains to LPS (Shi *et al*., 2014) or to bacteria through GBPs (Santos *et al*., 2020; Wandel *et al*., 2020) facilitates the multimerization of caspase-4 monomers, although how this process activates caspase-4 proteolytic function is undetermined. Homodimerisation of caspase-4 and -11 occurs through interactions within the catalytic domain (Fuentes-Prior & Salvesen, 2004). While dimerisation of caspase-11 is alone sufficient for it to acquire proteolytic activity (Ross *et al*., 2018), IDL autoproteolysis is additionally triggered to produce an active p32/p10 tetrameric species (Lee *et al*, 2018; Ross *et al*., 2018). Caspase-11 IDL processing to generate p32/p10 is required for caspase-11 to cleave GSDMD and induce pyroptosis. Caspase-11 does not appear to be able to self-cleave at the CDL to inactivate its protease function (Ross *et al*., 2018).

Caspase-4 cleavage fragments of approximately 20 kDa and 32 kDa containing the large protease subunit are reported upon non-canonical inflammasome signalling in human macrophages (Casson *et al*., 2015; Wang *et al*, 2020). However several aspects of caspase-4 cleavage are unknown, including their location (e.g. CDL versus IDL) and the responsible protease (caspase-4 itself versus another protease). Also undetermined is whether dimerisation and auto-processing affects caspase-4 activity and substrate specificity within the non-canonical inflammasome.

This study elucidates the molecular basis of caspase-4 activation during non-canonical inflammasome signalling. We demonstrate that caspase-4 needs to dimerise and be auto-processed at the IDL to generate fully active caspase-4 p34/p9 and p32/p9 protease species, which then cleave GSDMD to induce pyroptosis. Surprisingly, and in contrast to murine caspase-11, active caspase-4 can also directly cleave pro-IL-1β into its mature form.

## Results

### Dimerisation and IDL cleavage activates caspase-4

Both caspase-1 and -11 require proximity-induced dimerisation to become activated (Boucher *et al*., 2018; Ross *et al*., 2018). It is possible the same is true for caspase-4. Given that the LPS-binding CARD domain is necessary for LPS activation of caspase-4, we hypothesised that LPS binding clusters caspase-4 monomers to facilitate proximity-induced dimerisation of the caspase-4 enzymatic subunits. The caspase-4 dimer can then gain basal catalytic activity, sufficient for auto-cleavage. To test whether dimerisation is necessary and sufficient for caspase-4 activation, we used the DmrB dimerisation system. This system allows precise control of caspase dimerisation (Boucher *et al*., 2018) that is independent of LPS interactions with the caspase-4 CARD domain. ΔCARD-caspase-4 was expressed fused to an N-terminal V5-tagged DmrB domain (**fig. 1A**). We anticipated that adding the dimeriser drug, AP20187, would trigger wildtype caspase-4 protease activity and resultant cleavage at one or more sites (**fig. 1B**), and that these activities would be ablated by mutation of the catalytic cysteine (C258A). The DmrB-caspase-4 constructs were expressed in human embryonic kidney (HEK) 293T cells, and caspase activity was measured by monitoring fluorescence generated by cleavage of the Ac-WEHD-AFC peptidic substrate. Indeed, AP20187-induced dimerisation of wildtype caspase-4 triggered Ac-WEHD-AFC cleavage and auto-processing, and this was ablated by C258A mutation of the catalytic cysteine (**fig. 1C, D, E**). Immunoblotting with an antibody that recognises the caspase-4 large subunit showed that AP20187 induced cleavage of the full-length p43 into two smaller fragments, p34 and p32 (**fig. 1B, E**), which V5 western blot identified as products of IDL cleavage (**fig. 1B, E**). Together, these results suggest that dimerisation induces caspase-4 basal activity, IDL auto-processing and Ac-WEHD-AFC cleavage. While it is possible that caspase-4 forms multimers through CARD-CARD interactions as previously reported (Shi *et al*., 2014), our data indicate that caspase-4 dimers are alone sufficient for protease activity.

**Figure 1.**
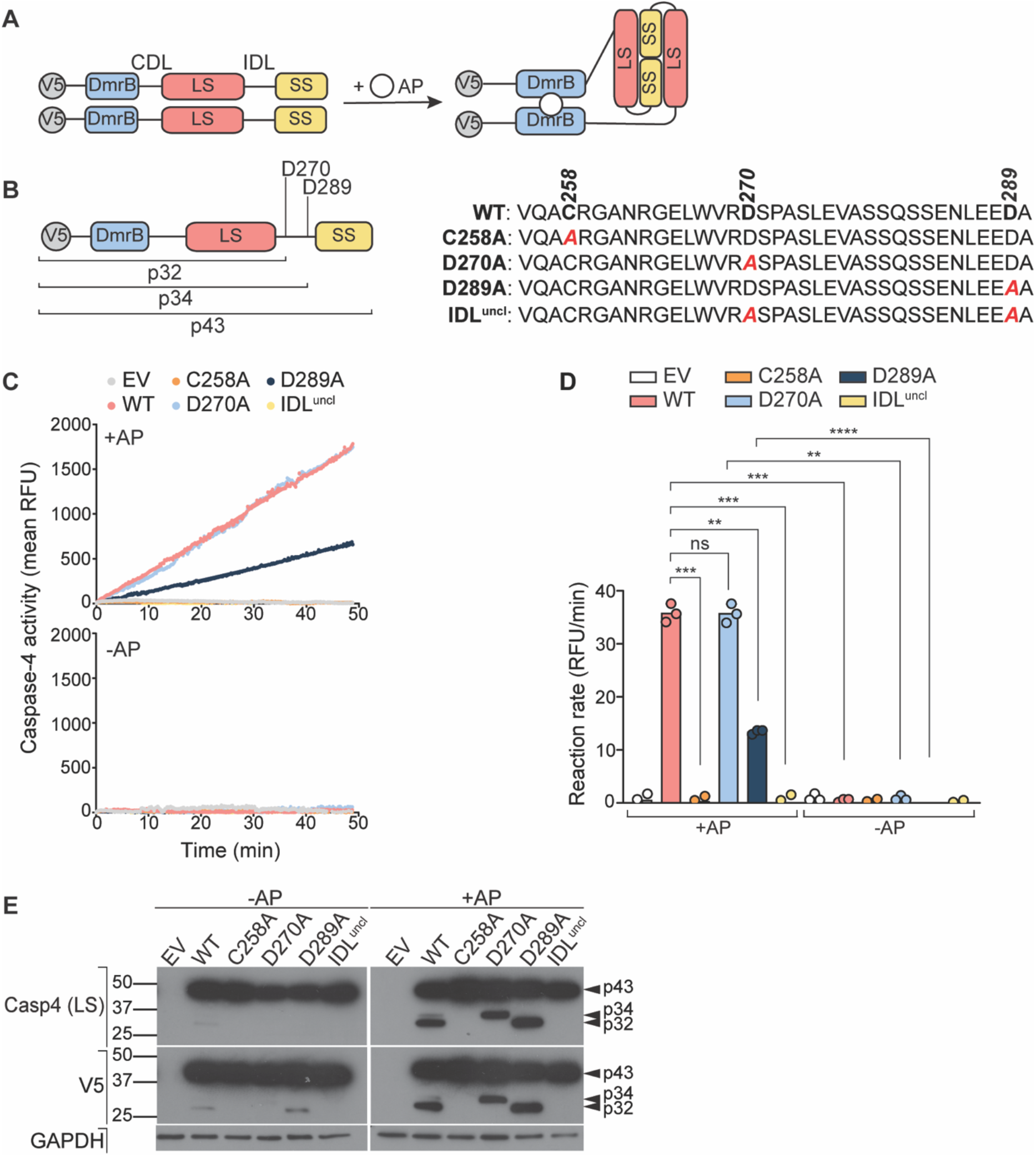
Dimerisation and IDL cleavage are required for caspase-4 proteolytic activity. (**A**) Schematic of caspase-4 dimerisation with the DmrB system, which allows controlled dimerisation by AP20187 (AP). All DmrB constructs were N-terminally V5-tagged. (**B**) Schematic of band sizes generated by cleavage at D270 and D289. These putative cleavage sites were mutated (D→A) to prevent auto-processing. An IDL^uncl^ construct encoding mutations at both D270 and D289 sites was also included. (**C**) Caspase-4 activity was measured by relative fluorescence (RFU) generated by Ac-WEHD-AFC substrate cleavage. Caspase-4 was expressed in HEK293T cells and incubated with AP for 30 min prior to Ac-WEHD-AFC cleavage experiment. Linear regression analysis (**D**) showing rate of Ac-WEHD-AFC cleavage. Data are mean ± SEM of three biological replicates. Each data point represents an individual donor. *p* ≤ 0.01 (**), *p* ≤ 0.001 (***), *p* ≤ 0.0001 (****). (**E**) Caspase-4 constructs were transfected in HEK293T cells and dimerised by AP. Auto-processing was analysed by western blot of cell extracts.

### Caspase-4 auto-processing at D270 and D289 is required for full protease activity against a peptide substrate

To first determine sites in the IDL that are cleaved during auto-processing, and the impact of their cleavage on caspase-4 function, we identified two putative cleavage sites, D270 (WVRD↓) and D289 (LEED↓) (**fig. 1B**), based on sequence similarity to the preferred cleavage sequence of (W/L)EHD↓ for caspase-4 (Kang *et al*, 2000; Thornberry *et al*, 1997). Two single-point mutations at D270 and D289, as well as a double mutant (IDL^uncl^), were created within the V5-DmrB-caspase-4 construct (**fig. 1B**) and tested for activity as before. Interestingly, the D270A single-point mutant retained full Ac-WEHD-AFC cleavage activity, while D289A mutation markedly diminished, although did not ablate, activity (**fig. 1C, D**). Western blots revealed that the generation of caspase-4 cleavage fragments required the catalytic cysteine and so were products of auto-celavage. p34 and p32 fragments corresponded to cleavage at D289 and D270, respectively, and that mutating one of these residues did not ablate processing at the other (**fig. 1E**). Importantly, mutation of both cleavage sites (IDL^uncl^) ablated caspase-4 protease activity on Ac-WEHD-AFC (**fig. 1C, D**) and prevented caspase-4 self-cleavage (**fig. 1E**). Together, these data indicate that IDL cleavage at either site (D289 or D270) is required for caspase-4 protease activity, with auto-processing at D289 required for full activity against the Ac-WEHD-AFC substrate. Interestingly, if cleavage at D289 is blocked by mutation, cleavage at D270 can partially compensate to generate a caspase-4 species with moderate activity. These results together indicate that dimerisation is both necessary and sufficient to induce caspase-4 auto-catalytic activity, IDL cleavage and activity against the Ac-WEHD-AFC substrate.

### Dimerisation and IDL auto-processing is required for caspase-4 to cleave GSDMD and IL-1β

We next sought to determine whether caspase-4 IDL auto-proteolysis is required for caspase-4 proteolytic activity against its natural substrate GSDMD. Given that IDL cleavage may expose recognition sites for caspase interaction with particular substrates (Liu *et al*, 2020; Wang *et al*., 2020), we also examined whether the site of IDL cleavage affects the repertoire of substrates processed by caspase-4 by co-expressing DmrB-caspase-4 constructs alongside V5-tagged caspase substrates (GSDMD and pro-IL-1β) in HEK293T cells. AP20187 was added to induce caspase-4 dimerisation and proteolytic activity, and cleavage of the substrates was examined. Wildtype caspase-4 cleaved GSDMD, while the IDL^uncl^ and the catalytic mutant did not (**fig. 2A**). While the D270 and D289 single mutants could cleave GSDMD, cleavage was dramatically reduced compared to wildtype caspase-4 (**fig. 2A**). This indicates that caspase-4 requires dimerisation and auto-processing at both IDL sites (D270 and D289) for efficient cleavage of GSDMD.

**Figure 2.**
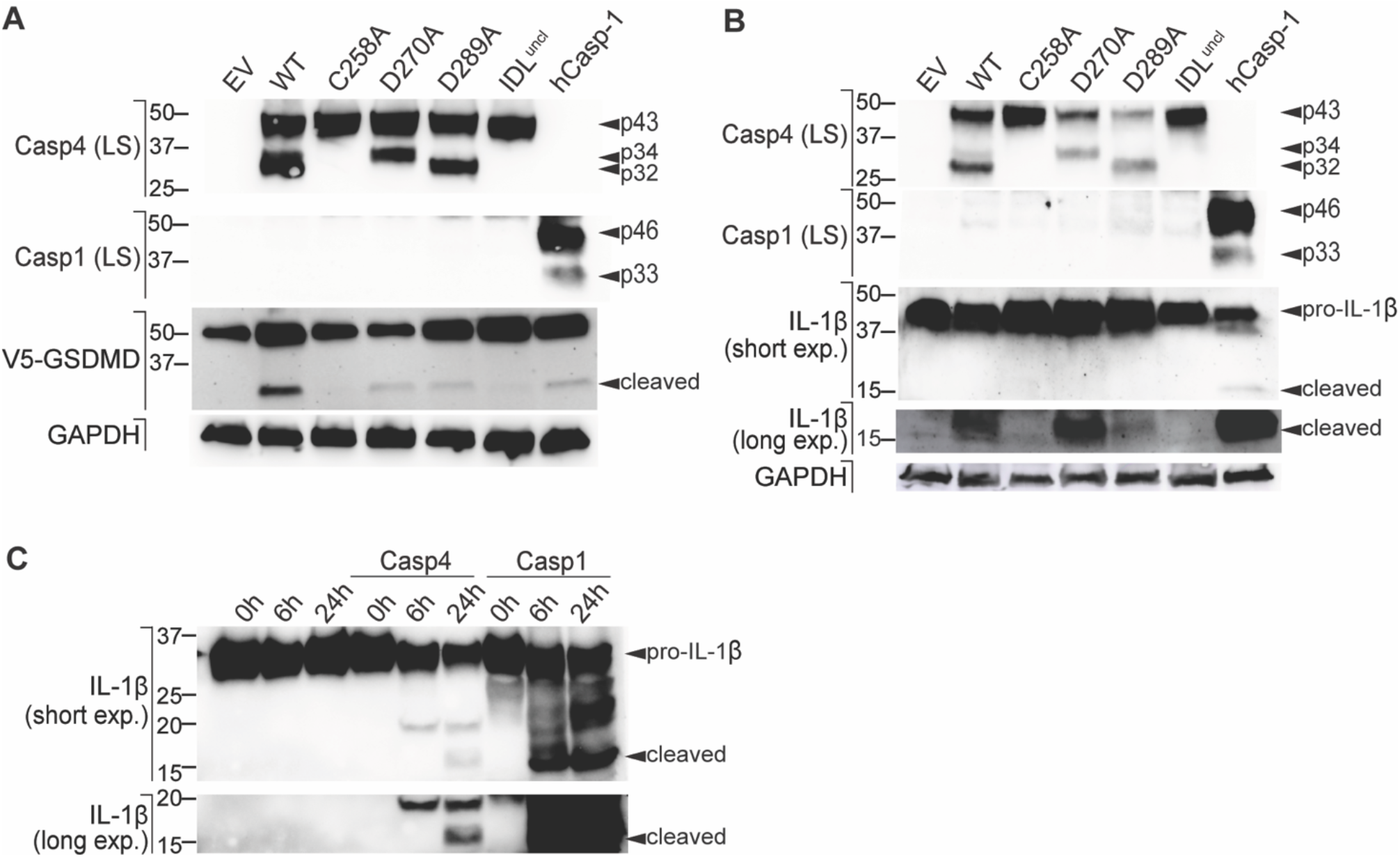
Dimerisation induces caspase-4 auto-processing and cleavage of GSDMD and IL-1β in cellulo and in vitro. (**A**-**B**) Caspase-4 constructs were co-expressed with (**A**) V5-GSDMD or (**B**) pro-IL-1β in HEK293T cells and caspase-4 was dimerised by cell exposure to AP20187 (AP). Substrate cleavage was analysed by western blot of the whole cell lysates. (**C**) Recombinant caspase-4 or caspase-1 was incubated with recombinant human pro-IL-1β for 0 h, 6 h, and 24 h. All data (**A**-**C**) are representative of three biological replicates.

Interestingly, dimerised caspase-4 was also able to cleave pro-IL-1β to its mature 17 kDa form. Pro-IL-1β was effectively cleaved by dimerised wildtype caspase-4 and the D270A single mutant, but not by the D289A single mutant, IDL^uncl^ or the catalytic mutant (**fig. 2B**). This suggests that caspase-4 can directly cleave pro-IL-1β if it is first dimerised and then self-cleaves at residue D289. Taken together, these data suggest that D270 cleavage is required for caspase-4 to efficiently cleave GSDMD, but not pro-IL-1β or Ac-WEHD-AFC (**fig. 1C, D**). Thus, caspase-4 self-cleavage at different IDL sites might allow for optimal recognition and processing of different substrates.

To confirm the unexpected finding that caspase-4 can directly cleave pro-IL-1β, we employed a fully recombinant system. Recombinant caspase-4 or human caspase-1 were incubated at 37°C with recombinant pro-IL-β for 0 h, 6 h, or 24 h. Here, caspase-4 cleaved pro-IL-1β to the bioactive 17 kDa form (**fig. 2C**), albeit at a much slower rate than caspase-1. Caspase-1 also processed pro-IL-1β to a 26 kDa fragment as previously reported (Afonina *et al*, 2015), while caspase-4 did not. Caspase-4 cleaved pro-IL-1β to a yet undefined fragment at ∼20 kDa by 6 h. Caspase-4 generated the mature 17 kDa form of IL-1β by by 24 h in this fully recombinant system, confirming caspase-4 as a bona fide IL-1β converting enzyme.

### Cytosolic LPS induces GSDMD and pro-IL-1β cleavage independently of the NLRP3 inflammasome

Transfected LPS is reported to interact directly with caspase-4 to induce caspase-4-driven death of human monocyte-derived macrophages (HMDMs), keratinocytes and epithelial cell lines (Casson *et al*., 2015; Knodler *et al*, 2014; Shi *et al*., 2014). Our earlier data indicates that caspase-4 directly cleaves GSDMD (**fig. 2A**), which is consistent with established functions for caspase-4 (and murine caspase-11) in initiating cell death independently of caspase-1. These data also suggested that caspase-4 directly cleaves pro-IL-1β, which is unexpected given that caspase-11 cannot. To investigate the potential for endogenous caspase-4 to directly cleave pro-IL-1β in a physiological setting, we primed HMDMs to upregulate pro-IL-1β expression, and then transfected the cells with *Escherichia coli* (*E. coli*) K12 LPS to activate the non-canonical caspase-4 inflammasome. For comparison, nigericin was added in parallel to stimulate caspase-1 activation by the canonical NLRP3 inflammasome, while primed HMDMs were also exposed to either the caspase-1/4 inhibitor VX-765 (Wannamaker *et al*, 2007) or the NLRP3-specific inhibitor MCC950 (Coll *et al*, 2019; Coll *et al*, 2022), before cell exposure to inflammasome activators. Inflammasome signalling was measured by LDH cytotoxicity assay for cell lysis, ELISA for IL-1β secretion and western blotting for cleaved caspase-1 and IL-1β. As expected, MCC950 suppressed nigericin-dependent cell death and IL-1β release, but not caspase-4-driven cell death induced by cytosolic LPS (**fig. 3A, B**). Importantly, LPS-induced secretion of mature IL-1β was minimally affected by MCC950 (**fig. 3A, C**), indicating that LPS-induced pro-IL-1β cleavage and release occurred independently of NLRP3 in human macrophages. The caspase-1/4 inhibitor VX-765 did suppress LPS-induced secretion of mature IL-1β (**fig. 3A, C**), confirming that this response was mediated by caspase-4. These results suggest that the non-canonical inflammasome signals via distinct mechanisms in human versus murine macrophages, and further support a molecular function for caspase-4 as an IL-1β-converting enzyme.

**Figure 3.**
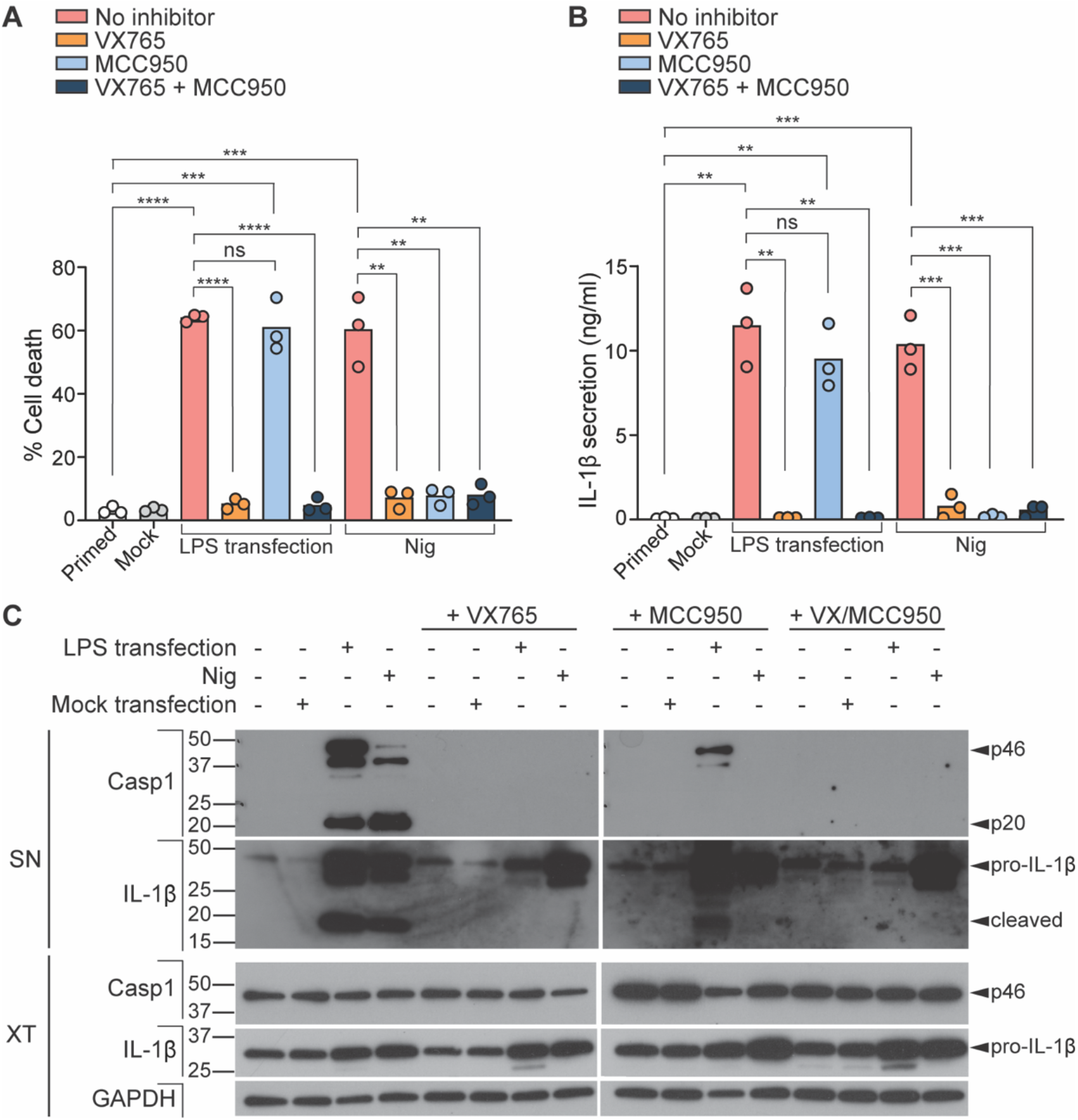
Cytosolic LPS induces GSDMD and IL-1β processing independently of NLRP3 or caspase-1. (**A**-**C**) HMDMs were primed with extracellular K12 LPS (100 ng/mL) for 4 h, then either transfected with K12 LPS (10 μg/mL) using Lipofectamine LTX transfection reagent or 10 μM nigericin (Nig) was added to wells. VX-765 (10 μM) and MCC950 (10 μM) was added to cells 1 h before LPS transfection or nigericin treatment, then 4 h later, supernatants and cell lysates were collected. (**A**) Secretion of mature IL-1β into the supernatant was assessed by ELISA. (**B**) Cell death was assessed by quantifying lactate dehydrogenase (LDH) release into the supernatant, compared with full lysis (Triton X-100) control. Data (**A**-**B**) are the mean ± SEM of three biological replicates, and significance was assessed by unpaired t-test. Each data point represents an individual donor. *p* ≤ 0.01 (**), *p* ≤ 0.001 (***), *p* ≤ 0.0001 (****). (**C**) HMDMs were analysed by western blot of precipitated supernatant (SN) or cell lysate (XT). Western blots are representative of three biological replicates.

### Cytosolic LPS induces cell death and IL-1β secretion in the absence of NLRP3

Caspase-4 is strongly expressed within epithelial barriers with important defence functions, such as bronchial epithelial cells and intestinal epithelial cells. To further investigate physiological functions for caspase-4 in IL-1β processing and secretion, we examined caspase-4 signalling in a human bronchial epithelial cell (HBEC-KT) cell line which does not express NLRP3 (**fig. 4A**). We primed HBEC-KT cells with LPS to upregulate pro-IL-1β expression, and then after priming, cells were exposed to caspase-1/4 (VX-765) and NLRP3 (MCC950) inhibitors. Cells were then transfected with *E. coli* K12 LPS to activate the caspase-4 inflammasome. LPS transfection induced HBEC-KT death and IL-1β secretion regardless of the presence of MCC950 (**fig. 4B, C**), confirming that NLRP3-activated caspase-1 was not responsible for either of these signalling outputs. The caspase-1/4 inhibitor VX-765 supressed LPS-induced caspase-4 cleavage (**fig. 4D**) and IL-1β secretion, confirming the involvement of active caspase-4 in IL-1β production.

**Figure 4.**
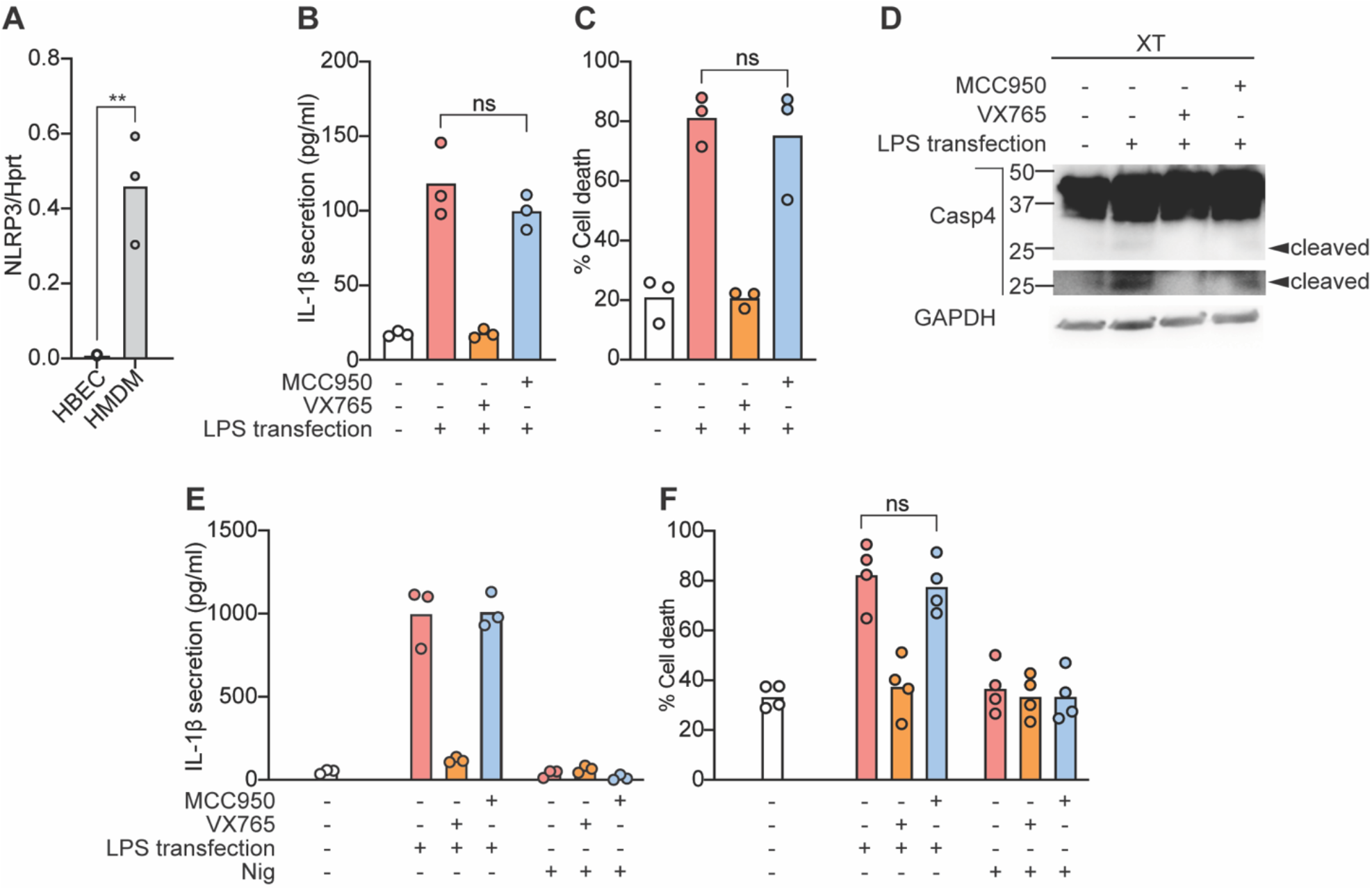
Cytosolic LPS induces cell death and NLRP3-independent IL-1β secretion. NLRP3 mRNA expression levels were compared between untreated HBEC-KT cells and HMDMs (**A**). HBEC-KT cells were primed with extracellular K12 LPS (100 ng/mL) for 4 h (**B-D**), or HIEC-6 cells were primed with IFNγ (10 ng/mL) for 16 h (**E-F**). The cells were then either transfected with K12 LPS (10 μg/mL) using Lipofectamine LTX transfection reagent or 10 μM nigericin (Nig) was added to wells. VX-765 (10 μM) or MCC950 (10 μM) was added to cells 1 h before LPS transfection or nigericin treatment. Cell supernatants and cell lysates were collected 4 (HMDM and HBEC-KT) or 6 h (HIEC-6) later. Secretion of mature IL-1β into the supernatant was assessed by ELISA (**B, E**). Cell death was assessed by quantifying lactate dehydrogenase (LDH) release into the supernatant, compared with full lysis (Triton X-100) control (**C, F**). HBEC-KT lysates were analysed by western blot and are representative of three biological replicates (**D**). Data in (**A-C, F**) are the mean ± SEM of three biological replicates, and significance was assessed by unpaired t-test. Each data point represents an individual donor. *p* ≤ 0.01 (**), *p* ≤ 0.001 (***), *p* ≤ 0.0001 (****). Data in (**E**) are mean and technical triplicate, and are representative of three biological replicates.

We further confirmed NLRP3-independent IL-1β secretion in a primary human fetal intestinal epithelial cell line (HIEC-6) which does not respond to the NLRP3 agonist, nigericin (**fig. 4E, F**). HIEC-6 were primed with IFNγ and transfected with LPS to activate caspase-4. Cytosolic LPS again induced IL-1β secretion in a manner insensitive to MCC950 but blocked by VX-765 (**fig. 4E, F**). These results collectively suggest that caspase-4 directly induces IL-1β production in HIEC-6 and HBEC-KT cells.

## Discussion

Innate immune responses are critical for host defence against invading pathogens, and thus require mechanisms for rapid microbial detection and response. The non-canonical inflammasome is a key signalling complex that recognises and responds to cytosolic Gram-negative bacteria. In humans, caspase-4 is constitutively expressed by many cells wherein it surveys the cytosol for bacterial LPS to sense Gram-negative bacterial infection (Knodler *et al*., 2014; Schmid-Burgk *et al*, 2015). In this study, we sought to determine the activation mechanisms of caspase-4 during non-canonical inflammasome signalling.

Previous studies suggested that LPS binding to the caspase-4 CARD domain results in caspase-4 clustering and multimerisation (Shi *et al*., 2014). By using the DmrB system to precisely control caspase dimerisation independently of CARD-mediated multimerisation, we showed that dimerisation alone is sufficient to confer basal activity; higher-order-complexes such as oligomers are thus dispensable. This is in support of the long-standing proximity-induced dimerisation model for initiator caspases and suggests that dimerisation of the catalytic subunits is the likely initial activation step for caspase-4. We further propose that caspase-4 CARD interactions with LPS bring caspase-4 monomer catalytic subunits into close proximity within the noncanonical inflammasome, thus enabling caspase-4 dimerisation and activation that is sufficient for caspase-4 IDL auto-cleavage.

We identified two cleavage sites within the IDL that modulate caspase-4 activity. Using the DmrB dimerisation system, we demonstrated that the IDL is cleaved at D270 and D289 to produce p32 and p34 fragments respectively. During the preparation of this manuscript, another report also identified these IDL cleavage sites (Wang *et al*., 2020), reporting that D289 cleavage is necessary for LPS-induced caspase-4 signalling and GSDMD processing, whereas D270 is partially dispensable. Our results further indicate that cleavage at D270 and D289 bestow different degrees of caspase-4 activity and substrate selectivity. Cleavage at either site conferred partial activity on GSDMD, whereas cleavage at both sites was required for optimal GSDMD processing. The finding that IDL cleavage is required for full caspase-4 activity on substrates suggests that cleavage at both IDL linker sites liberates an exosite for substrate recognition. In fact, the cleavage sites D270 and D289 are flanked by two important residues – W267 and V291 – for hydrophobic interaction with GSDMD (Wang *et al*., 2020). Cleavage at both D270 and D289 is thus likely to unmask the exosite for efficient GSDMD processing.

We identified pro-IL-1β as a novel substrate of caspase-4. Earlier studies suggested that recombinant caspase-4 may cleave mouse IL-1β in vitro (Devant *et al*, 2021; Kamens *et al*, 1995), but until now, caspase-4 is generally considered to be unable to cleave human pro-IL-1β (Bibo-Verdugo *et al*, 2020; Devant *et al*., 2021; Downs *et al*, 2020). Here, we demonstrate that caspase-4 directly cleaves human IL-1β in both recombinant and cellular systems. Using the DmrB dimerisation system, we showed that IDL D289 cleavage is required for pro-IL-1β processing in cells, whereas D270 appears to be dispensable, suggesting that pro-IL-1β is cleaved by the caspase-4 species p34/9, and perhaps p32/p9. Further, purified recombinant caspase-4 cleaved purified recombinant human pro-IL-1β. Using two different human epithelial cell lines, we further show that caspase-4 is responsible for NLRP3-independent IL-β production in fully endogenous cellular systems. NLRP3 expression is generally restricted to the myeloid compartment (Guarda *et al*, 2011), while caspase-4 is expressed more broadly (e.g. at barrier surfaces that regularly encounter bacteria). The capacity for caspase-4 to initiate both pyroptosis and IL-1β maturation could be particularly important in non-myeloid cells, such as epithelial cells, that are unable to signal via NLRP3. The IL-1β converting enzyme activity of caspase-4 thus allows these cells to mount strong and rapid pro-inflammatory responses at a site of barrier compromise, thereby preventing bacterial dissemination.

This study found that the activation mechanisms of human caspase-4 has several parallels and differences to the elucidated mechanisms of murine caspase-11. For both caspases, dimerisation of catalytic subunits appears to be the initial step for achieving proteolytic activity. Caspase-4 and caspase-11 both acquire basal protease activity upon dimerisation, while higher-order complexes such as multimers are dispensable. Our data further suggest that the activation mechanisms and molecular functions of caspase-4 differ from caspase-11 in two significant ways. Firstly, caspase-4 facilitates NLRP3-independent IL-1β production by directly cleaving human pro-IL-1β into its mature, active form. Secondly, two auto-processing sites within the caspase-4 IDL enable the potential generation of at least four different active dimeric caspase-4 species with distinct functions: (i) full length p43, which can auto-process, (ii) D270-cleaved p32/p11, which has moderate GSDMD-cleavage activity, (iii) D289-cleaved p34/p9, which has moderate GSDMD-cleavage activity and IL-1β convertase activity, and (iv) D270/D289-cleaved p32/p9, which has strong GSDMD-cleavage activity. This differs to caspase-11, which harbours only one IDL cleavage site (Ross *et al*., 2018), and which cannot process IL-1β. Similarly, pro-IL18 has been found to be a substrate for caspase-4 but not caspase-11 (Santos *et al*., 2020; Wandel *et al*., 2020), suggesting that the functions of pro-inflammatory caspases have diverged across evolution.

In summary, this study presents the first detailed mechanistic model for caspase-4 activation and its function on the non-canonical inflammasome. The mechanistic insight into caspase-4 activation pathways described herein delivers new understanding of the molecular processes controlling protective responses during infection, as well as the pathological inflammatory responses driven by non-canonical inflammasome signalling.

## Methods

### Cell lines

All cells were cultured in humidified incubators at 37°C and with 5% CO_2_. HEK293T (ATCC CRL-3216) cells were cultured in Dulbecco’s modified Eagle’s medium (DMEM; Gibco) supplemented with 10% heat-inactivated fetal bovine serum (FBS) and 1% penicillin-streptomycin (Pen-Strep). HEK293T cells were used to express the DmrB constructs and were seeded in 10 cm cell culture dishes. The DrmB constructs were cloned into pEF6 vectors (Invitrogen) and then transfected into HEK293T cells using lipofectamine 2000 (Thermo Fisher) transfection reagent. For *in cellulo* substrate cleavage experiments, substrate-containing plasmids (V5-GSDMD, IL-1β) were co-transfected into HEK293T cells alongside caspase vectors. After overnight incubation at 37°C, the cells were re-seeded at a density of 1 × 10^6^ cells/mL. HBEC-KT (ATCC CRL-4051) cells were cultured in Airway Epithelial Cell Basal Medium (AECBM; ATCC) supplemented with Bronchial Epithelial Cell Growth Kit (ATCC). HBEC-KT cells were seeded at a density of 1 × 10^6^ cells/mL and primed for 4 h with extracellular ultrapure *Escherichia coli* K12 LPS (100 ng/mL) before inflammasome activation. HIEC-6 (ATCC CRL-3266) cells were cultured in reduced serum media (Opti-MEM; Gobco) supplemented with 4% FBS, 1% Pen-Strep, 1xGlutaMAX, 10 ng/mL of Epithelial Growth Factor (EGF), and 20 mM HEPES (Gibco). HIEC-6 cells were primed for 16 h with interferon-gamma (10 ng/mL) before inflammasome activation.

### Primary cell culture

Studies using primary human cells were approved by the UQ Human Medical Research Ethics Committee. The Australian Red Cross Blood Service provided buffy coats from anonymous, informed and consenting adults for this research study. Human peripheral blood mononuclear cells were isolated from screened buffy coats by density centrifugation with Ficoll-Plaque Plus (GE Healthcare) followed by magnetic-assisted cell sorting, according to standard protocols (Schroder *et al*, 2012). Monocyte-derived macrophages were differentiated with recombinant human CSF-1 (150 ng/mL; endotoxin-free, produced in insect cells by the UQ Protein Expression Facility) at 37°C with 5% CO_2_. On day 6 of differentiation, HMDMs were plated at a density of 800,000 cells/mL in Roswell Park Memorial Institute (RPMI; Gibco) 1640 media supplemented with 10% heat-inactivated fetal bovine serum, 1% penicillin-streptomycin, 1x GlutaMAX and 150 ng/mL CSF-1. HMDMs were used for experimentation on day 7 of their differentiation. HMDMs were primed for 4 h with extracellular ultrapure *Escherichia coli* K12 LPS (100 ng/mL) before inflammasome activation.

### DmrB caspase-4 dimerisation and substrate co-expression cleavage assays

The medium from transfected HEK293T cells was replaced with Opti-MEM containing 1 μM of B/B Homodimeriser (AP20187; Clontech) and incubated for 30 min. The medium was replaced with caspase activity buffer (200 mM NaCl, 50 mM HEPES, pH 8.0, 50 mM KCl) supplemented with 100 μg/mL digitonin, 10 mM DTT, and 100 μM Ac-WEHD-AFC. Processing of Ac-WEHD-AFC was monitored at 37°C for regular time intervals using the M1000 TECAN spectrofluorometer (400 nm excitation, 505 nm emission). Cell extracts and supernatants were precipitated using methanol/chloroform and analysed by western blot following previously described methods (Groß, 2012) and the following reagents: antibodies against the caspase-4 large subunit, hIL-1β, caspase-1 large subunit (Bally-1, mouse monoclonal, 1:1000; Adipogen), V5 (SV5-Pk1, mouse monoclonal, 1:2000; Abcam) and GAPDH.

### In vitro recombinant protein cleavage assays

5 U of full-length human caspase-1 (Abcam) or caspase-4 (Abcam) recombinant proteins were incubated with 2 μg of recombinant human pro-IL-1β (Sino Biological Inc) in caspase activity buffer (200 mM NaCl, 50 mM HEPES, pH 8.0, 50 mM KCl). The protein mix was incubated at 37°C for 0 h, 6 h, and 24 h. Cleavage of recombinant proteins was analysed by western blot using standard methods (Groß, 2012) and the following reagents: antibodies against the caspase-4 large subunit (4B9, mouse monoclonal antibody, 1:1000; Santa Cruz Biotechnology), human IL-1β (goat polyclonal antibody, 1:1000, R&D Systems), caspase-1 (D7F10, rabbit monoclonal, 1:1000; Cell Signalling Technology).

### Inflammasome assays

To activate the non-canonical inflammasome, cells were transfected with ultrapure *Escherichia coli* K12 LPS (10 μg/mL) using 0.25% Lipofectamine LTX (Thermo Fisher) and following the manufacturer’s instructions. To activate the NLRP3 canonical inflammasome, 10 μM nigericin sodium salt (Sigma-Aldrich) was added for 2 or 4 h. Where cells were exposed to 10 μM MCC950 or 10 μM VX-765 (MedChemExpress), these were added 1 h prior to the inflammasome agonists by replacing medium with Opti-MEM replete with inhibitors. IL-1β secretion into the supernatant was assessed by ELISA (R&D systems), according to the manufacturer’s instructions. Cell cytotoxicity was measured using the CytoTox96 Non-radioactive Cytotoxicity Assay (Promega) and expressed as a percentage of total cellular LDH (100% lysis control). Cell extracts and methanol/chloroform-precipitated supernatants were analysed by western blot using standard methods (Groß, 2012) and the following reagents: antibodies against the caspase-4 large subunit (4B9, mouse monoclonal antibody, 1:1000; Santa Cruz Biotechnology), human IL-1β (goat polyclonal antibody, 1:1000, R&D Systems), caspase-1 (D7F10, rabbit monoclonal, 1:1000; Cell Signalling Technology), GSDMD (rabbit polyclonal antibody, 1:1000; Cusabio) and GAPDH (polyclonal rabbit antibody, 1:5000; BioScientific).

### mRNA expression analysis

Cell monolayers were harvested in 350 μL Buffer RLT with b-mercaptoethanol, and RNA was processed using the RNeasy Mini Kit (Qiagen) with on-column DNA digestion following the manufacturer’s protocol. RNA concentrations were determined on NanoDrop spectrophotometer and an equal amount of RNA from each sample were used for downstream cDNA synthesis. cDNAs were synthesized by reverse transcription using Superscript III Reverse Transcriptase (ThermoFisher) with OligoDT priming. Gene expression was quantified in 384 well plates by qPCR using SYBR green reagent (Applied Biosystems) on a QuantStudio 7 Flex Real-Time PCR System (ThermoFisher). Relative gene expression was determined using the change-in-threshold (2^-DDCT^) method with Hypoxanthine Phosphoribosyltransferase 1 (HPRT) as an endogenous control.

### Statistical analysis

Statistical analysis was performed using GraphPad Prism 8.0 software. Biological replicates were pooled by combining the means of technical replicates. Data were analysed for normality using the Shapiro-Wilk normality test. For caspase activity assays, Ac-WEHD-AFC activity curves were analysed by linear regression to determine the slope (relative fluorescence units/min) then tested for statistical significance using parametric paired *t* tests (two-sided). Data were considered significant when *p* ≤ 0.05 (*), *p* ≤ 0.005 (**), *p* ≤ 0.0005 (***), and *p* ≤ 0.0001 (****). Data from ELISA and LDH assays were tested for statistical significance using parametric unpaired *t* tests (two-sided). Data were considered significant when *p* ≤ 0.01 (*), *p* ≤ 0.001 (**), *p* ≤ 0.0001 (****).

## Acknowledgements

We gratefully acknowledge Dr. Fiona Wylie for editing this manuscript. This work was supported by the Australian Research Council (Discovery Project DP190102285), the National Health and Medical Research Council of Australia (Fellowships 1141131 and 2009075 to KS; Synergy Grant 2009677 to KS), The University of Queensland (Postdoctoral Fellowship to DB), and a Springboard Award from the Academy of Medical Science and the Wellcome Trust (SBF006/1025 to DB). KS is a co-inventor on patent applications for NLRP3 inhibitors licensed to Inflazome Ltd, a company headquartered in Dublin, Ireland. Inflazome is developing drugs that target the NLRP3 inflammasome to address unmet clinical needs in inflammatory disease. KS served on the Scientific Advisory Board of Inflazome in 2016–2017, and serves as a consultant to Quench Bio, USA and Novartis, Switzerland. The authors have no additional financial interests.

## Author Contributions

AHC, LIL, DB, and KS designed the experiments. AHC, XW, DB, PV, and JvP executed and analysed experiments. D. Boucher and K. Schroder designed the project and supervised the study. AHC and KS wrote the paper with input from all authors.

